# Cell-free genetic devices confer autonomic and adaptive properties to hydrogels

**DOI:** 10.1101/2019.12.12.872622

**Authors:** Colette J. Whitfield, Alice M. Banks, Gema Dura, John Love, Jonathan E. Fieldsend, Sarah A. Goodchild, David A. Fulton, Thomas P. Howard

**Affiliations:** School of Natural and Environmental Sciences, Faculty of Science, Agriculture and Engineering, Newcastle University, Newcastle-upon-Tyne, NE1 7RU, U. K.; Biosciences, College of Life and Environmental Sciences, University of Exeter, Exeter, EX4 4PY, U.K.; Computer Science, College of Engineering, Mathematics and Physical Sciences, University of Exeter, EX4 4PY, U. K.; Defence Science and Technology Laboratory, Porton Down, Wiltshire, SP4 0JQ, U. K.

**Keywords:** Cell-free Protein Synthesis, Synthetic Biology, Hydrogels

## Abstract

Smart materials are able to alter one or more of their properties in response to defined stimuli. Our ability to design and create such materials, however, does not match the diversity and specificity of responses seen within the biological domain. We propose that relocation of molecular phenomena from living cells into hydrogels can be used to confer smart functionality to materials. We establish that cell-free protein synthesis can be conducted in agarose hydrogels, that gene expression occurs throughout the material and that co-expression of genes is possible. We demonstrate that gene expression can be controlled transcriptionally (using *in gel* gene interactions) and translationally in response to small molecule and nucleic acid triggers. We use this system to design and build a genetic device that can alter the structural property of its chassis material in response to exogenous stimuli. Importantly, we establish that a wide range of hydrogels are appropriate chassis for cell-free synthetic biology, meaning a designer may alter both the genetic and hydrogel components according to the requirements of a given application. We probe the relationship between the physical structure of the gel and *in gel* protein synthesis and reveal that the material itself may act as a macromolecular crowder enhancing protein synthesis. Given the extensive range of genetically encoded information processing networks in the living kingdom and the structural and chemical diversity of hydrogels, this work establishes a model by which cell-free synthetic biology can be used to create autonomic and adaptive materials.

**Significance statement:** Smart materials have the ability to change one or more of their properties (e.g. structure, shape or function) in response to specific triggers. They have applications ranging from light-sensitive sunglasses and drug delivery systems to shape-memory alloys and self-healing coatings. The ability to programme such materials, however, is basic compared to the ability of a living organism to observe, understand and respond to its environment. Here we demonstrate the relocation of biological information processing systems from cells to materials. We achieved this by operating small, programmable genetic devices outside the confines of a living cell and inside hydrogel matrices. These results establish a method for developing materials functionally enhanced with molecular machinery from biological systems.

## Introduction

Cell-free protein synthesis (CFPS), uses cellular transcriptional and translational machinery to synthesise proteins outside the living cell (1). CFPS systems have shown potential as diagnostic tools in the detection of Zika and Ebola viruses (2, 3), as well as for detection of metabolic molecules such as hippuric acid and cocaine (4). CFPS is typically performed in liquid reactions *in vitro*, though cell-free expression of reporter gene constructs has been demonstrated from gene networks preserved on paper (3) and in the presence of agarose (5), clay (6) DNA (7, 8), fibrin and PEG-peptide (9) hydrogels. Importantly, hydrogels have valuable properties in their own right. They act as pastes (10), lubricants (11) and adhesives (12) or in the provision of structural support for regenerative cell growth (13). Considering the applications of hydrogels and the functionality of CFPS we hypothesised that hydrogel matrices may act as a physical chassis for CFPS – allowing their move from an *in vitro* to a physical application – while simultaneously augmenting the function of the hydrogel.

Smart materials are materials capable of changing one or more of their properties in a predictable and useful manner in response to defined stimuli (14, 15). These capabilities are integral to the material itself. They may be described as autonomic – meaning they sense, diagnose and respond to external stimuli – and adaptive – meaning they can reconfigure their functionality, structure or shape. Current applications of smart materials include rapid light-responsive photochromic inks in sun-glass (16, 17), fabric sensors that detect external stimuli (18), soft robotic micromachines (19, 20) and drug delivery systems (21, 22). Smart hydrogels have been developed that change physical properties in response to alterations in pH, ionic strength or temperature (23, 24) while advanced actuation responses (25) and logical information processing capabilities (26) have also been demonstrated. Our ability to design and build autonomic and adaptive materials, however, remains basic compared to the multi-responsive, multi-functional properties observed within the biological domain. In this regard, we have much to learn from living systems, where local molecular and biochemical reactions form a hierarchical organisation from which highly complex behaviours emerge.

The field of molecular biomimetics provides opportunities to repurpose molecular and biochemical functionality for the development of new smart materials (27). To date, however, effort has largely centred on the bottom-up construction of novel, bio-inspired or bio-constructed materials (28-30). Here, rather than using molecular tools to construct materials, we instead transpose the molecular machinery for protein synthesis from *Escherichia coli* into hydrogel matrices. This permits the cell-free expression of genes in a physical chassis that are subsequently used to augment the hydrogel with autonomic and adaptive properties. We show that both expression and co-expression of proteins occurs in materials operating without an external liquid. We established sensory functionality in agarose gels regulated at both the transcriptional and translational level, and we built on this through the regulated expression of enzyme activity that altered the physical structure of the chassis in response to an exogenous trigger molecule. Importantly, we find that CFPS is compatible with a wide range of hydrogels and that even in gels in which protein synthesis is hampered, the expression of functional enzymes that act on the chassis material result in changes to macroscopic mechanical properties. These findings demonstrate that we can use cell-free genetic devices to confer autonomic and adaptive properties to ostensibly unresponsive materials. As such, it becomes possible to tap directly into the diversity and specificity of biological sensing and information processing networks to create novel, smart materials.

## Results

### Establishing agarose as a chassis for cell-free protein synthesis

Cell-free synthesis of single reporter proteins has been demonstrated in the presence of hydrogels previously (5-9). In each instance however, CFPS reactions have either been assembled in liquid form prior to mixing with hydrogels (5), hydrogels are bathed or submerged in CFPS reaction mixtures (7, 8), or hydrogel polymers form around a liquid CFPS reaction (9). As such we cannot exclude the possibility that CFPS reactions are occurring as liquid reactions and protein products are diffusing into the gel or alternatively that protein synthesis is occurring before the gel has formed. If CFPS is to be used to engineer autonomic or adaptive capabilities into hydrogels, it is critical that CFPS occurs throughout the material, not simply at the interface between the liquid and solid phases, or with minor entry into the gel. We first sought to establish agarose as an appropriate model for testing our proposal that embedding CFPS into hydrogels can be used to confer smart functionality to materials. We established single gene expression of mCherry and eGFP in agarose gels (Fig. 1A). The gels were prepared by the addition of molten 3 % agarose to cell-free reaction components in the appropriate cast. Gels were allowed to set for 10 min at room temperature before incubation at 37 °C to ensure all reaction components were situated within the hydrogel polymer network. There is no external liquid phase in our system. Confocal microscopy was then used to image at 25 μm intervals throughout the gel. Our data indicated that mCherry- and eGFP-fluorescence was located evenly throughout the material (Fig. 1B, SI Appendix Fig. S1). It is critical to our goals that such materials are able to support the synthesis of more than a single gene product and to our knowledge, previous studies have only sought to express a single gene product in the presence of hydrogels. We therefore confirmed co-expression of two fluorescent proteins in an individual gel chassis. We monitored the expression of both mCherry and eGFP in agarose using confocal microscopy and observed that both proteins were expressed with no spatial competition between proteins (Fig. 1C). An overlay of both the red and green wavelength ranges visualise the even spatial distribution of both proteins. Agarose is therefore capable of supporting CFPS as a stand-alone system in the absence of an external liquid phase. Agarose gels containing only a single template (mCherry or eGFP) were also prepared and then brought into contact with each other. Again, expression of the correct protein is observed in the appropriate chassis, but it was also observed that there is signal diffusion between the neighbouring gels (Fig. 1D). This illustrates a key advantage of deploying cell-free devices in gels - that gene expression functionality can be spatially organised in a manner that is not possible in liquid phase reactions.

**Fig. 1.**
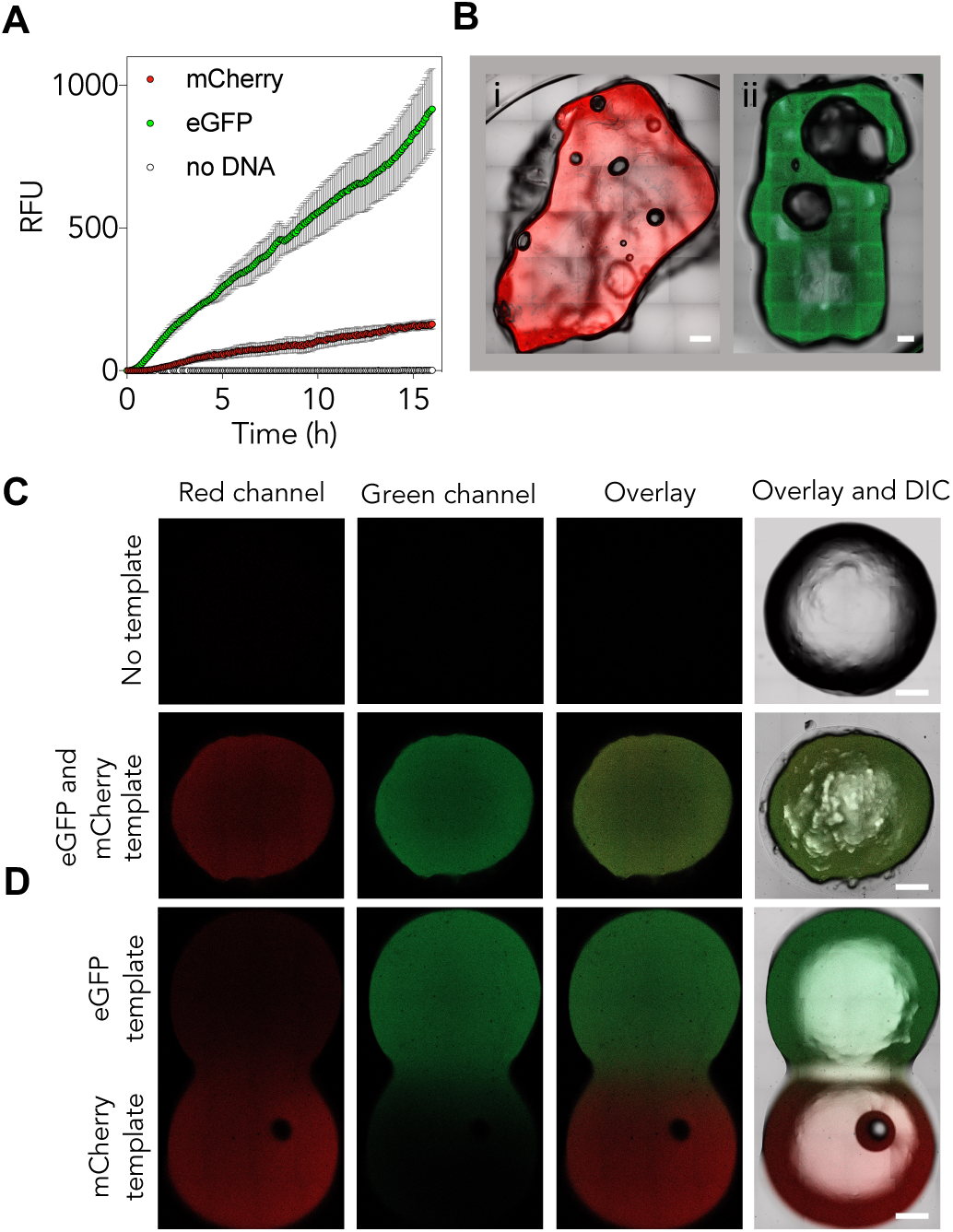
Fluorescent protein expression within an agarose chassis. (*A*) Expression of mCherry and eGFP is possible in 0.75 % agarose. Fluorescence was monitored over time (*n* = 3, error bars are SE mean). (*B*) Expression of mCherry (i) and eGFP (ii) visible throughout the gel. Confocal microscopy of a central plane shows fluorescence of both mCherry and eGFP in each hydrogel. (*C*) Co-expression of both mCherry and eGFP is possible in a single agarose chassis. 0.75 % agarose gels were prepared with CFPS reagents, incubated for 4 h and imaged in the red and green channels. Also shown are the two channels overlaid, and the differential interference contrast (DIC) overlaid with both channels. Gels either contained 4 µg of each template or lacked DNA template altogether. (*D*) Spatial organisation of gene expression in agarose. Two 0.75 % agarose gels were prepared separately with 4 µg of either eGFP or mCherry template in each. The gels were placed into contact with one another and incubated for 4 h before imaging. Scale bars = 1 mm.

### Cell-free genetic devices operating in an agarose chassis can be designed for autonomic and adaptive responses

To generate autonomic features (i.e. stimuli-responses) in materials it is important to demonstrate that exogenous signals can be used to manipulate gene expression within a gel. We explored this at both the transcriptional and translational regulatory level. First, we used cell-free extract from *E. coli* BL21(DE3) cells that possess an endogenously expressed *lac* repressor protein (LacR) and placed mCherry expression under the control of the LacR-repressed *trc* promoter (31) (Fig. 2A). Binding of the LacR protein to the *trc* promoter should supress mCherry production and be relieved by the addition of isopropylthio-β-galactoside (IPTG) which binds to the LacR protein and in turn prevents it repressing the *trc* promoter. As expected, when this construct was included in the gel, the addition of IPTG resulted in a 1.3-fold increase in mCherry fluorescence (Fig. 2A and 2D). To test whether transcriptional repression of gene expression by a cell-free synthesised transcriptional regulator is possible, we performed CFPS in *E. coli* Rosetta2 cell-free extract (Fig. 2B). Rosetta2 cells lack the LacR regulator protein. In this instance a LacR construct under control of the constitutive promoter J23100 was incubated in the agarose-based CFPS system for 2 h to permit accumulation of the repressor protein prior to addition of the *trc*:mCherry construct either in the presence or absence of IPTG. We observe that gene expression is repressed in the absence of IPTG, and relieved in the presence of IPTG (Fig. 2B). To further improve the system response, we introduced the construct encoding LacR to the *E. coli* BL21(DE3) cell lysates, providing both a pre-made and CFPS derived LacR repressor. In this instance, the addition of the LacR template further enhanced *trc* repression and could be relieved by the addition of IPTG providing 3.4-fold greater fluorescence production than the repressed sample (Fig. 2C-D).

**Fig. 2.**
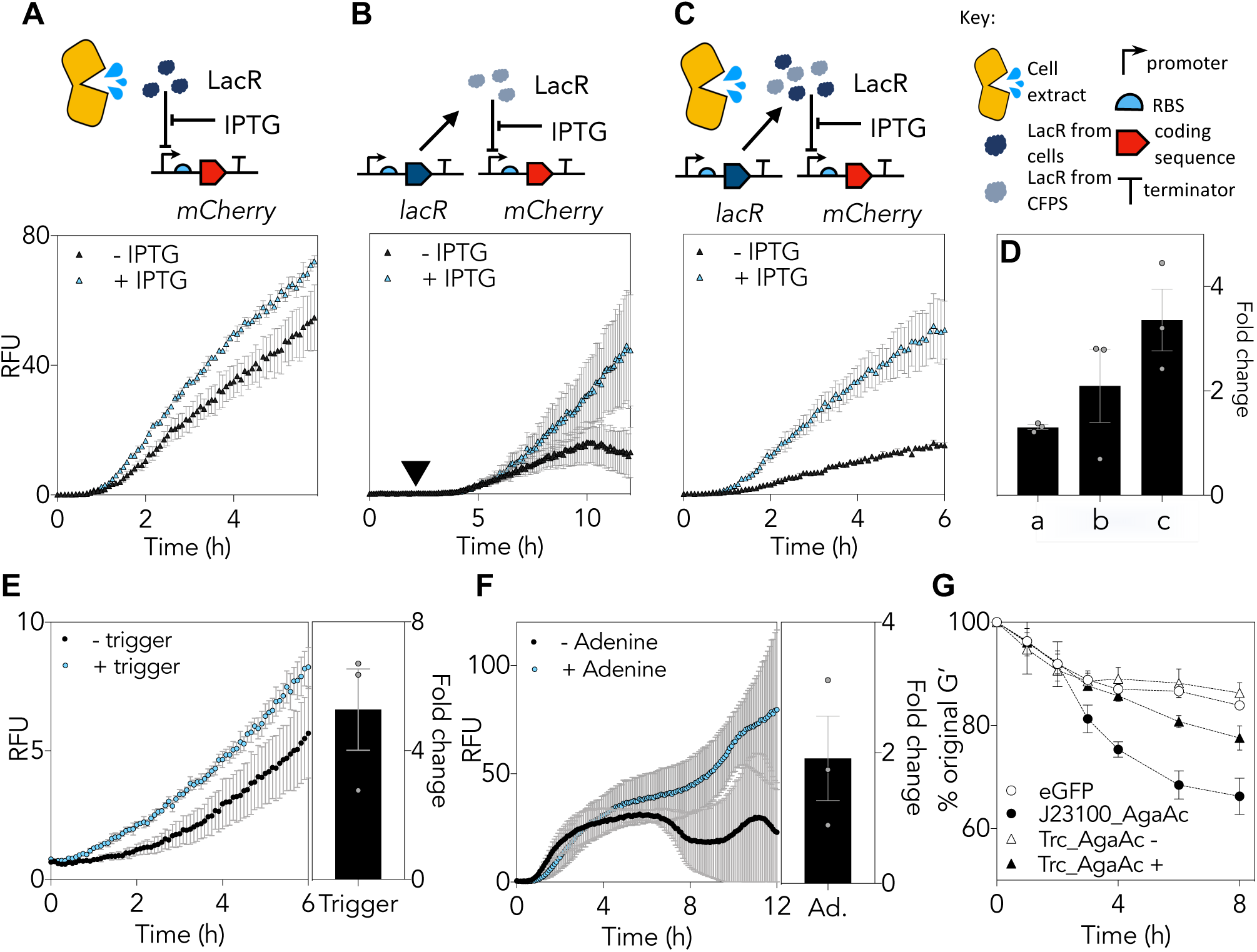
Regulating genetic devices within an agarose chassis can stimulate alterations in the physical properties. (*A*) mCherry synthesis under control of the trc promoter in BL21(DE3) cell-free extract containing endogenous LacR protein either in the presence (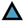) or absence (▲) of 40 µM IPTG. (*B*) mCherry synthesis in Rosetta2 cell-free extract lacking endogenous LacR protein. The *lacR* template under the J23100 promoter was allowed to incubate for 2 hours before the addition of trc:*mCherry* template in the presence of (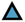) or absence of (▲) 10 µM IPTG. (*C*) mCherry synthesis under control of the trc promoter in BL21(DE3) cell-free extract containing endogenous LacR protein and *lacR* template under control of the 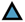 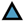 J23100 promoter, either in the presence () or absence (▲) of 40 µM IPTG. (*D*) The fold change between the presence and absence of IPTG in A-C. (*E*) and (*F*) Cell-free translational control utilising Riboswitches; (*E*), expression of GFP under the control of a toehold switch in the presence and absence of the trigger RNA template, (*F*) expression of GFP under the control of an adenine responsive riboswitch in the presence and absence of 0.5 µM adenine. (*G*) Rheological analysis monitoring the % change decrease of the stiffness, G’, of 0.75 % agarose in the presence of three DNA constructs in the same experimental conditions as **c** and in the absence (Δ) and presence of IPTG (▲) (*n* = 3, errors bars are SE mean).

A second means of regulating gene expression is at the translational level. Under these circumstances, DNA is transcribed to mRNA, but its translation through to the corresponding polypeptide is inhibited. One mechanism for achieving this is through the use of riboswitches (32, 33). Riboswitches sequester important sequence information (such as ribosome binding sites and start codons) into mRNA secondary structure folds. When these RNA switches bind their cognate ligand their conformation alters, exposing the hidden regulatory elements thereby permitting gene expression. Establishing riboswitch functions in gels broadens the mechanisms for molecule detection further and, because translational control responses are potentially quicker than transcriptional mechanisms, provides routes to faster material responses. In the first instance, we used a toehold device which responds to a 77-nucleotide ligand (34). The results show that presence of the riboswitch at the 5’ end of the transcript inhibits CFPS of GFP while in the presence of the trigger ligand there is a 5.3-fold increase in GFP fluorescence over a 6 h period (Fig. 2E). Next we demonstrated that a gel-based riboswitch was able to respond to the small molecule adenine. The adenine-responsive riboswitch, *addA* has previously been described in a cell-based assay (35), but not in cell-free systems. Again, we observed the expected repression of GFP fluorescence that was relieved almost 2-fold in the presence of adenine (Fig. 2F). Taken together these data demonstrate that gene expression in gels can be moderated by both transcriptional and translational control mechanisms. Unlike conventional chemo-responsive materials that rely on trigger molecule reactivity, this establishes responses to compounds with low inherent chemical reactivity

Finally, one of the clear benefits of operating cell-free gene networks in a physical chassis is the potential to allow the sensory input of the genetic device to translate into an output response that alters the structure or function of the material itself. We therefore examined whether it was possible to alter the physical structure of an agarose chassis through the cell-free synthesis of the agarose enzyme, AgaAc (36). In the first instance, expression of AgaAc was under control of the constitutive promoter J23100. We monitored cell-free digestion by following the change in the hydrogel’s storage modulus (G’) by rheology. We found that constitutive expression of an agarase led to a 21 % reduction in G’ in comparison to gels expressing only eGFP (Fig. 2G). Next we placed the AgaAc coding sequence downstream of the *trc* promoter and - using an *E. coli* BL21(DE3) cell-free extract - observed the effect of IPTG on the system. We observed a reduction of 10 % in G’ of the gel in the presence of IPTG compared to gels without IPTG, which behaved in a manner equivalent to control reactions (Fig. 2G). The results establish that cell-free operation of genetic devices can be used to convert the ostensibly unreactive hydrogel agarose into a stimuli-responsive smart material.

### Hydrogels as chassis for cell-free synthetic biology

Hydrogels have a vast range of physical properties and functionalities which we may wish to combine with the capabilities offered by cell-free genetic devices. To ascertain whether CFPS is broadly compatible with a range of hydrogels we expressed a single fluorescent mCherry reporter protein in 12 gels with a variety of different structural and chemical characteristics, including entangled polymers, micellar aggregates and covalently crosslinked materials. Two fabrication methods were required to integrate the cellular components within the hydrogels (Fig. 3A): in Method A, the gel was prepared, freeze-dried and reconstituted with the CFPS reaction components; in Method B, the CFPS reaction components were combined, freeze-dried, and re-constituted with liquid hydrogel. For some systems, both methods were appropriate, whilst for others, only one method was feasible (SI Appendix Table S1 and SI Appendix Fig. S2). Two controls were employed: a positive control (CFPS of mCherry in liquid phase) and a negative control (CFPS without DNA template ensuring that all fluorescence detected could be attributed to mCherry protein synthesis). Our data indicated that the ability to perform CFPS in hydrogels, though variable, is widespread (Fig. 3B, SI Appendix Table S1 and SI Appendix Fig. S2). Of the 12 gels tested, six were polysaccharide-based hydrogels: agarose, agar, xanthan gum, hyaluronic acid, Gelzan™ and alginate. These gels feature extensive noncovalent linkages between polymer chains, formed by hydrogen bonding and Van der Waals forces (agarose, agar, xanthan and hyaluronic acid) or Mg^2+^ or Ca^2+^-enabled ionic bridges (for Gelzan™ and alginate respectively). For agarose, agar, xanthan, and Gelzan™ the fluorescence detected was equivalent to, or up to 400 % greater than, the fluorescence observed when CFPS was conducted in liquid phase. For alginate and hyaluronic acid in contrast, there was a reduction in fluorescence of between 13 and 57 % respectively compared to the control. Three peptide-based hydrogels were also assessed; collagen, gelatin and HydroMatrix™. These gels are formed through hydrophobic interactions, π-π stacking, hydrogen bonding and electrostatic interactions between polymers (37). For each of the three peptide-based hydrogels CFPS was observed above baseline but signals were low, reaching between 6 and 43 % of that observed in the positive control. F-108 and F-127 are poloxamer gels where individual micelles aggregate to form a gel. For these gels, fluorescence was equivalent to, or up to, 150 % of the liquid phase control. Finally, an examination of CFPS in a polyacrylamide gel, a covalently crosslinked network of acrylamide and bis-acrylamide, revealed that CFPS-fluorescence was 130 % of that observed in the liquid phase control reaction. CFPS is therefore possible in a range of hydrogel materials and, in some instances, performing the reaction in a hydrogel chassis increases the fluorescent output of the reaction. Importantly, it follows that choice of hydrogel chassis for CFPS is not restrictive, and that a gel can be selected that is appropriate to the proposed end-use.

**Fig. 3.**
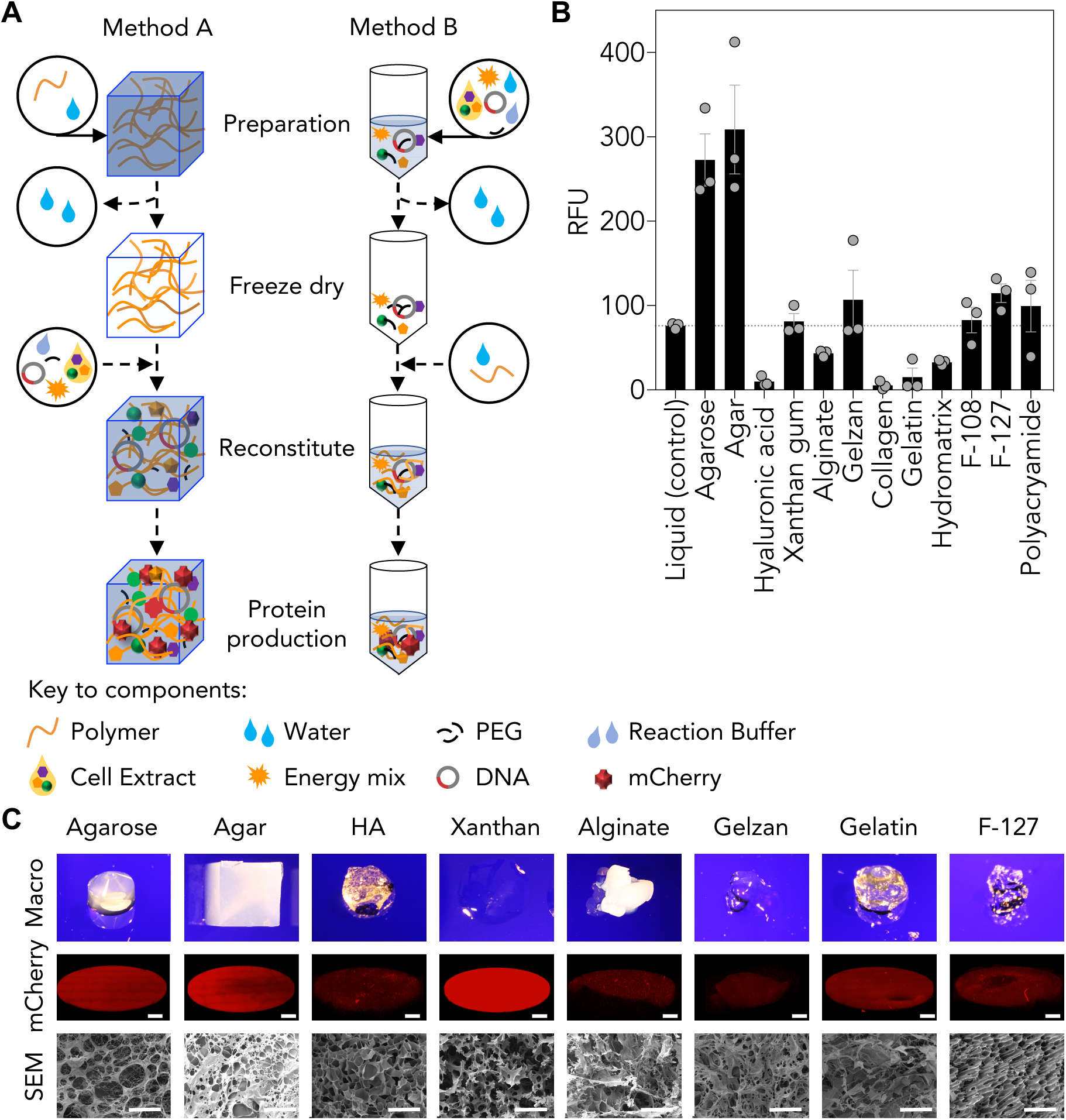
Cell-free gene expression of mCherry within hydrogel chassis. (*A*) Cell-free protein synthesis of the mCherry fluorescent reporter follows one of two methods. Method A requires the initial preparation of materials, followed by freeze-drying the material and reconstitution with the cell-free reagents. Method B requires the initial preparation of the cell-free reagents, followed by freeze-drying and reconstitution with liquid hydrogel. (*B*) CFPS is possible in a range of materials. Relative fluorescence units (RFU) detected from a range of hydrogels containing cell-free reaction mixtures and mCherry template DNA. RFU is represented as the max RFU detected over a 16 h period. Hydrogel % ^w^/_v_ plotted; agarose: 1 %, agar: 1 %, hyaluronic acid: 5 %, xanthan gum: 0.5 %, alginate: 5 %, Gelzan™: 2 %, collagen: 1 %, gelatin: 10 %, HydroMatrix™: 1 %, F-108: 30 %, F-127: 30 %, polyacrylamide: 5 % (*n* = 3, error bars are standard error (SE) mean). (*C*) The hydrogels selected exhibit a range of macro- and micro-scale structures. Hydrogel images of the macro-structure (top), confocal fluorescence microscopy of a central plane of the hydrogels after 4 h of incubation at 37 °C with the CF components and mCherry coding DNA (middle, scale bar = 1 mm) and Scanning Electron Microscopy (SEM) of each material (bottom, scale bar = 100 µm).

As previously, it was important to confirm that gene expression occurs throughout the material. Using confocal microscopy, we again observed mCherry fluorescence throughout the gels (Fig. 3C and SI Appendix Fig. S3). Gels demonstrating equivalent or increased fluorescence compared to control reactions (agarose, agar, xanthan, gelatin and the poloxamer gels) displayed homogenous mCherry-fluorescence throughout the gel. Others, in which cell-free production of mCherry was detected but reduced (e.g. hyaluronic acid) demonstrated heterogenous patterns of fluorescence. With this in mind, we examined the physical properties of each hydrogel. Examination of a selection of gels using SEM (Fig. 3C) suggests that large, open pores correspond with homogenous patterns of gene expression than those with smaller, or more irregular shapes pores. This hypothesis is support by data which shows that mCherry fluorescence is greater in gels possessing high diffusion and water content (SI Appendix Figs. S4-S8, SI Appendix Tables S2-S3). The physical structure of each gel therefore plays a key role influencing cell-free protein synthesis

### The physical structure of the chassis affects cell-free protein synthesis, with diffusion and molecular crowding important factors in performance

To examine more directly how the physical nature of the hydrogel effects CFPS we returned to our model agarose system. We manipulated the matrix concentration of agarose and found that altering the concentration of the agarose matrix markedly alters the porosity of the gel and the diffusion of the dye fluorescein through the gel (Fig. 4A-C). The data also indicates that gene expression broadly correlates positively with fluorescein diffusion (Fig. 4D (blue)). The results, however, also indicate that there are non-linear responses to changes in matrix concentration indicating that diffusion is not the only factor that influences gene expression in agarose. We hypothesised that a further factor that could modify the observed responses is if the hydrogel network can also act as a macromolecular crowder. Macromolecular crowding is known to positively influence CFPS reactions (38, 39). We therefore additionally assayed CFPS without the crowding agent polyethylene glycol (PEG) and with PEG at half the standard concentration. Reactions lacking PEG were observed to have reductions in CFPS-dependent fluorescence of 72 % and those at half standard concentrations have reductions of 42 % (Fig. 4D inset). Interestingly, the data indicated that the gel matrix itself mitigates against the removal of PEG (Fig. 4D). Agarose gels of between 1 – 2 % ^w^/_v_ that lack PEG maintained fluorescence values equivalent to those observed in standard liquid phase reactions supporting the hypothesis that the hydrogel itself is acting as a structural crowding agent.

**Fig. 4.**
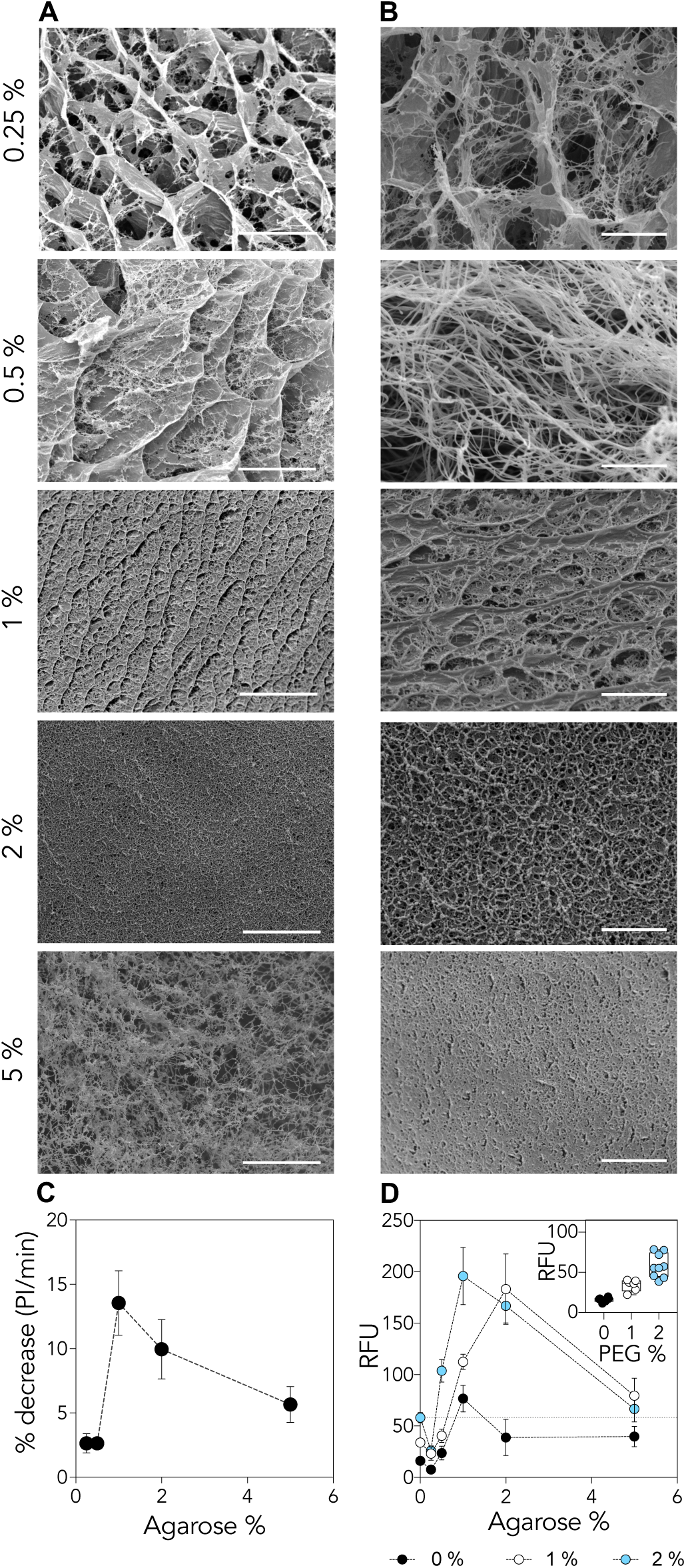
Physical impacts of a solid matrix on gene expression. (*A*), (*B*) SEM images of 0.25, 0.5, 1, 2 and 5 % agarose hydrogels. Scale bars (*A*) 20 µm and (*B*) 5 µm. (*C*) Analysis of fluorescein diffusion through agarose gels. Data was collected using a fluorescence microscope and intensity was determined using ImageJ. Data is represented as a % decrease over 5 min in fluorescence intensity (*n* = 3, error bars are SE mean). (*D*) Cell-free mCherry protein synthesis at increasing concentrations of agarose in the absence (•), or in the presence of 1 % (•) or 2 % (•) polyethylene glycol (PEG) (*n* ≥ 6, error bars are SE mean). The horizontal dashed line indicates performance of the standard liquid phase reaction in the presence of 2 % PEG. The insert graph demonstrates the effect of crowding on standard liquid phase CFPS.

### Cell-free protein synthesis of enzymes that target the chassis can result in large-differences in material structure

The appearance of mCherry fluorescence in hyaluronic acid was found to be much reduced compared to controls but was nonetheless reproducibly present. We wanted to probe whether, even in this scenario, expression of an enzyme targeting the material structure would result in a detectable physical outcome. We therefore placed a polysaccharide Family 7 lyase from *Klebsiella pneumoniae* capable of digesting hyaluronic acid (40) downstream of the constitutive promoter J23100 (Fig. 5A) and monitored cell-free digestion by following the change in G’ by rheology and swelling ability (Fig. 5B, and SI Appendix Fig. S9). A 25 % reduction in the G’ of a hyaluronic acid gel was observed over 4 h of lyase CFPS compared to a control reaction synthesising eGFP (Fig. 6B). To improve performance, we synthesised a chemically crosslinked hyaluronic acid : 1,4-butanediol diglycidyl ether (HA:BDDE) hydrogel (41) with more robust physical characteristics (Fig. 5C). CFPS of mCherry in HA:BDDE was confirmed and seen to be comparable to that observed in HA (SI Appendix Fig. S10) while the new, cross-linked hyaluronic acid gel was seen to have greater structural stability over the course of 8 h (Fig. 5D). Finally, digestion of HA:BDDE by a CFPS-expressed lyase was confirmed by rheology and swelling (Fig. 5D and SI Appendix Fig. S11). A reduction of one third in G’ of the HA:BDDE gel was seen compared to control. Interestingly, changes in the size and colour of the hydrogel could also be observed by eye (Fig. 5E and SI Appendix Fig. S12). These structural changes are particularly noteworthy as expression of proteins within hyaluronic acid gels is low compared to other materials (Fig. 3B). Even in this scenario, substantial adaptive changes in the material chassis can be instigated by expressing functional enzymes.

**Fig. 5.**
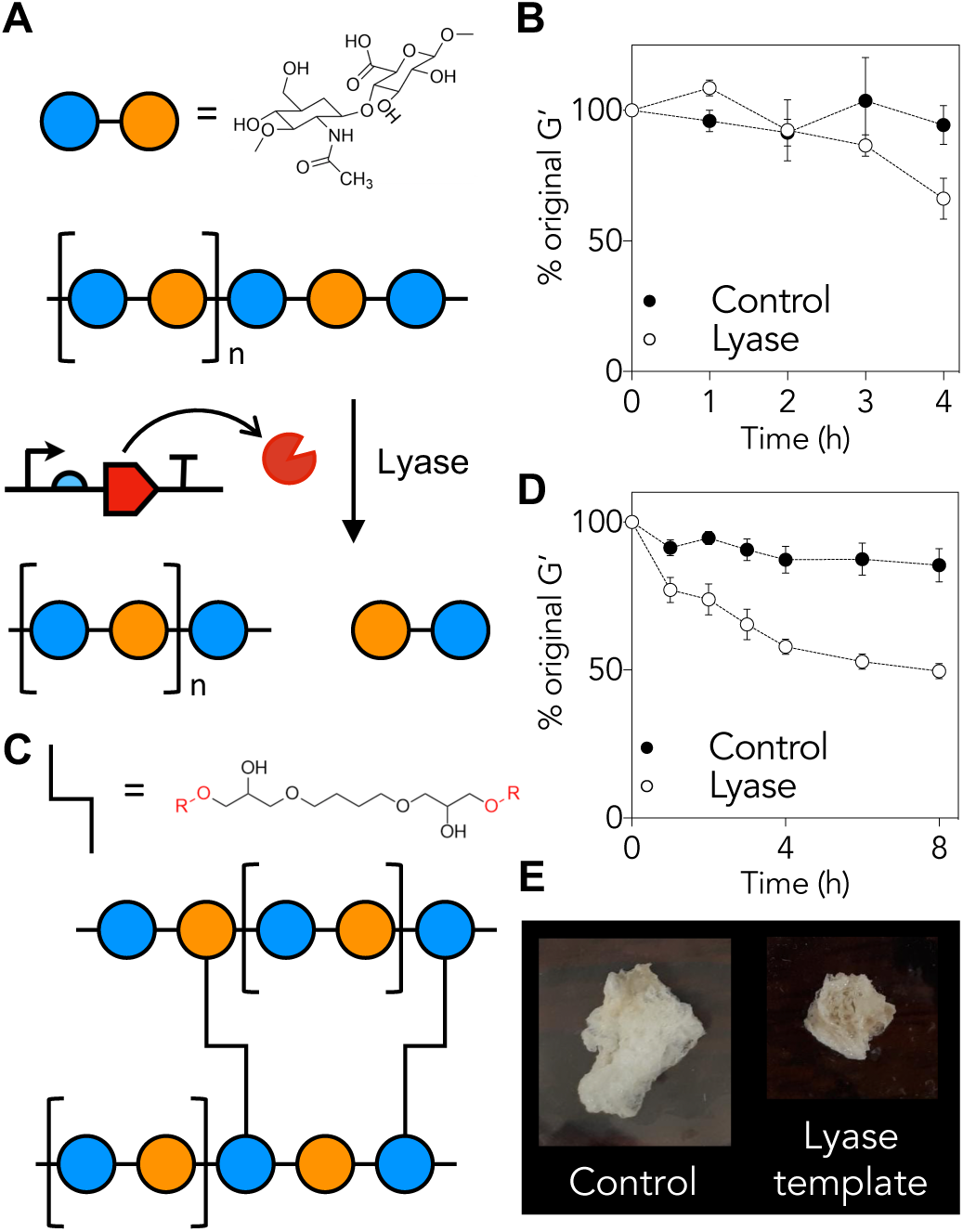
Genetic devices can stimulate alterations in the physical properties of hydrogel chassis. (*A*) Schematic of the digestion of hyaluronic acid (HA) (poly(β-glucuronic acid-[1→3]-β-N-acetylglucosamine-[1→4]) by a cell-free synthesised lyase and the chemical crosslink of 1,4-butanediol diglycidyl ether (BDDE) to HA. (*B*) Rheological analysis monitoring the % change decrease of the stiffness, G□, of HA over time in the presence of cell-free reagents with lyase template (•) and with eGFP template (•) (*n* = 3, errors bars are SE mean). (*C*) Rheological analysis monitoring the % change decrease of the stiffness, G□, of HA:BDDE over time in the presence of cell-free reagents with lyase template (•) and with eGFP template (•) (*n* = 5, errors bars are SE mean). (*D*) Visual image of gels after 48 h incubation in the absence of and with the lyase template. (*E*) Rheological analysis monitoring the % change decrease of the stiffness, G’, of 0.75 % agarose in the presence of three DNA constructs and in the absence (△) and presence of IPTG (▲) (*n* = 3, errors bars are SE mean).

## Discussion

We hypothesised that the combination of hydrogel functionality with the information processing systems embedded in cellular machinery could provide routes to developing new autonomic and adaptive materials. We have shown that hydrogel chassis permit expression of at least two genes within the material as well as the control of gene expression at both the transcriptional and translational level, demonstrating autonomic functionality. In contrast to conventional chemo-responsive materials that require reactive molecular triggers, for example, oxidants, reductants, acids or bases, here we have demonstrated the use of trigger molecules which possess low inherent chemical reactivity. Given the range of chemistries detected by biological systems coupled with increasing capabilities in designing synthetic gene networks, this work establishes a clear path to developing autonomic sensory systems within materials based upon molecular biology tools.

We have shown too that a wide range of hydrogels, including polysaccharide, proteinaceous and micellar gels, can house CFPS reactions. This is key - it follows that the choice of hydrogel chassis for these bioactivated materials is broad, not narrow, and that the choice of hydrogel can be apposite to its desired use. Finally, the expression of enzymes within the chassis that alter its structural properties demonstrates a route to establishing adaptive functionality. Together, these findings support our hypothesis that it is possible to design and build materials containing aspects of biological function by transposing cellular and molecular phenomenon from living systems into non-living chassis. Though cell-free synthetic biology remains technically challenging there are reports of the implementation of increasingly ambitious, programmable synthetic gene networks in cell-free environments (42-44), as well as a growing repertoire of transcriptional and translational ‘bioparts’ becoming available. These include small molecule (4, 45), nucleic acid (2, 3) and optogenetic (46) parts for the regulation of gene expression. Implementing genetic circuitry in hydrogels will allow their deployment in a physical and material manner that is not currently possible. Given the varied nature and commercial application of hydrogels, such devices could be used to enhance the sensory and adaptive functionality of existing infrastructure retrospectively, or for them to be seamlessly embedded in newly manufactured or imagined materials.

## Materials and Methods

### Materials

All materials were purchased from Sigma Aldrich (unless stated otherwise) except; oligos ordered from Eurofins, synthetic gene fragments from Integrated DNA Technologies (IDT) and the EcoFlex kit (47) from Addgene.

### Molecular Cloning

The pTU1-A-RFP backbone from the EcoFlex kit was amplified to remove the RFP coding region using 0.5 µM of primers (SI Appendix Table S4), 0.2 mM dNTPs, 5 ng plasmid, 1 × Q5 reaction buffer (New England Biolabs (NEB)) and 1 U of Q5 DNA polymerase in a 50 μL total volume. The amplification solution was heated to 98 °C for 30 s, 98 °C for 10 s and 72 °C for 70 s for 30 cycles before cooling to 4 °C. Purification was performed using a QIAquick PCR purification kit (QIAGEN).

EcoFlex (47) cloning followed the following procedure: 10 ng of PCR fragment from pTU1-A-RFP was combined with 10 ng of pBP-J23100, pBP-pET-RBS, pBP_eGFP, pBP_mCherry or the synthetic gene fragment (fragments were designed with EcoFlex compatible ends) and pBP-Bba_B0015 with 2 mg mL^-1^ bovine serum albumin, 10 U BsaI, 1 × T4 DNA ligase buffer and 3 U T4 DNA ligase. The combined solution was made up to 15 μL with H_2_O and cycled 15 × between 37 °C 10 for 5 min and 16 °C for 10 min before heat inactivation at 50 °C for 10 min and 80 °C for 10 min. The reaction mix was stored at 4 °C until the transformation step.

The EcoFlex reaction product was transformed into One Shot TOP10 chemically competent *E. coli* cells (ThermoFisher) following the manufactures protocol. 250 μL transformation solution was spread onto 100 µgmL^-1^ ampicillin selection LB agar plates and incubated at 37 °C overnight. A single colony was picked, grown over night in 5 mL LB and 100 μg mL^-1^ ampicillin and DNA were purified using a QIAprep Spin Miniprep Kit (QIAGEN). Sequencing was performed by GATC using the primers in SI Appendix Table S4. Following sequence verification, plasmid purification was performed using a QIAGEN Plasmid Maxi Kit following manufacturers protocol. Construct details are available in SI Appendix Table S5.

### Hydrogel preparation

Hydrogels were prepared following one of two methods (Fig. 1A). For Method A, hydrogel volumes may be different to the final required reaction volume (see SI Appendix Table S2 for details). To freeze dry hydrogels or reagents, samples were placed at −80 °C for over 30 min then transferred directly to the freeze dryer. Hydrogels can also be prepared fresh with CF components. Detailed descriptions of hydrogel preparations are provided in SI Appendix Table S6.

### Cell-free protein synthesis

Cell-free protein synthesis reaction composition followed the method described in ‘Hydrogel preparation’ and adding the CF components; 10 μL CF extract (final concentration of 8.9 mg/mL protein prepared following (48)), 25 μL 2 × energy buffer (final concentration 4.5 mM-10.5 mM Mg-glutamate, 40-160 mM K-glutamate, 0.33-3.33 mM DTT, 1.5 mM each amino acid except leucine, 1.25 mM leucine, 50 mM HEPES, 1.5 mM ATP and GTP, 0.9 mM CTP and UTP, 0.2 mg/ml tRNA, 0.26 mM CoA, 0.33 mM NAD, 0.75 mM cAMP, 0.068 mM folinic acid, 1 mM spermidine, 30 mM 3-PGA, 2 % PEG-8000 (unless stated otherwise)), 4 μg plasmid DNA (except for regulatory plasmids) assembled to 50 μL final volume using nanopure H_2_O. mCherry fluorescence was measured at 587 nm excitation and 610 nm emission while eGFP fluorescence was measured at 488 nm excitation and 512 nm emission using a Varioskan LUX multimode reader (ThermoFisher). All reactions were performed at 37 °C.

### Transcriptional and translational regulation

Transcriptional regulation of CFPS was performed with 4 μg of repression *lac* plasmid and 1 μg trc responsive plasmid. Translational regulation was performed with 8 μg trigger plasmid and 1 μg responsive plasmid for the toehold system and 1 μg template for the adenine responsive system (see SI Appendix Methods and SI Appendix Table S5 for further details).

### Confocal microscopy

A Zeiss LSM800 Airyscan/Spinning Disk with a DIC (differential interference contrast) was used for confocal microscopy with a 5 x magnification lens. For mCherry analysis a 561 nm laser was utilised and fluorescence was detected between 610 and 700 nm. For eGFP analysis, a 488 nm laser for eGFP and detect up to 520 nm.

### Material Characterisation

#### Fluorescein diffusion

Fluorescein was used to assess relative diffusion between each hydrogel at the best performing CFPS % ^w^/_v_. 10 μL of the liquid hydrogels was placed on a glass slide and once set, 0.2 μL 50 μM fluorescein in 1 × PBS (phosphate-buffered saline) was placed at the centre of the gel. Fluorescein diffusion was monitored using a Leica DM6-B Microscope (Leica, U.K.) with a DFC9000GT camera and a Platinum Bright 495 filter and exposed for 1 s per image (*n* = 3). The fluorescein intensity was measured at the centre of the original fluorescein drop site using ImageJ (49). Diffusion (D) was calculated following equation (1).

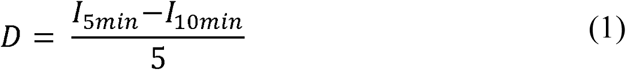

#### Rheology

Rheological tests were carried out on a HR-2 Discovery Hybrid Rheometer from TA Instruments equipped with a temperature controller. Rheological experiments were performed in 20 mm parallel-plate geometry using 500 μL of hydrogels (resulting in a gap size of 1.5 mm). The strain and the frequency were set to 1 % and 1 Hz, respectively. For rheological testing of gels expressing proteins, hydrogels were prepared to 500 μL (10 × volumes) in a Petri dish.

#### Water content

200 μL of material was prepared to the desired ^w^/_v_ ratio, followed by weighing the material, freeze drying and then re-weighing the dried matrix. The water content of each hydrogel was determined by calculating the % difference between the fresh and freeze-dried material. Each material at each ^w^/_v_ ratio was performed in triplicate.

#### Swelling capacity

500 μL of each material was prepared in a Petri dish, allowed to set and weighed to gain initial weights (m_i_). The gels were then submerged in H_2_O for 2 h, followed by removing the water and reweighing to determine the swollen weight of the hydrogel (m_s_). The swelling ratio (S) was determined following equation (2). Each hydrogel ^w^/_v_ was analysed in triplicate.

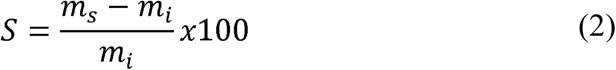

#### Hydrogel reconstitution

The level of reconstitution that each gel can undergo was determined by; preparing 200 μL of each gel in a 1.5 mL microcentrifuge tube and weighing the combined weight of gel and tube (m_i_), lyophilising the gel overnight and reweighing (m_l_) followed by the addition of 200 μL H_2_O. After 30 min, any unabsorbed H_2_O was removed followed by reweighing. The % reconstitution (R) is determined following equation (3). Each hydrogel ^w^/_v_ was analysed in triplicate.

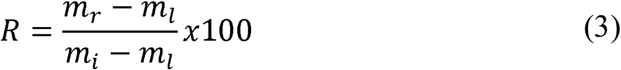

## Acknowledgements

This research is supported by the Engineering and Physical Sciences Research Council – Defence Science and Technology Laboratories award EP/N026683/1 (CJW, AB, JL, JEF, DAF, TPH) and the Biotechnology and Biological Sciences Research Council award BB/M018318/1 (GD, DAF). The authors wish to thank Dr Neil Dixon (BB/ R000069/1 and BB/K014773/1) and Mr Ross Kent (BB/M011208/1), University of Manchester, UK, for the kind gift of the *add-A*_GFP construct.

## Additional Information

### SI Appendix

accompanies this paper.

### Competing interests

The authors declare no competing interests.

