## Supplemental Information for "Cell-free genetic devices confer autonomic and adaptive properties to hydrogels"

**Supplementary Figures**

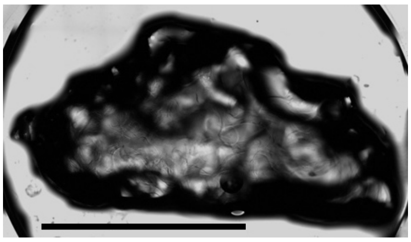

**Fig. S1** Confocal microscopy of 0.75 % agarose prepared with cell-free components after 4 h of incubation at 37 °C in the absence of template DNA. Scale bar is 10 mm.

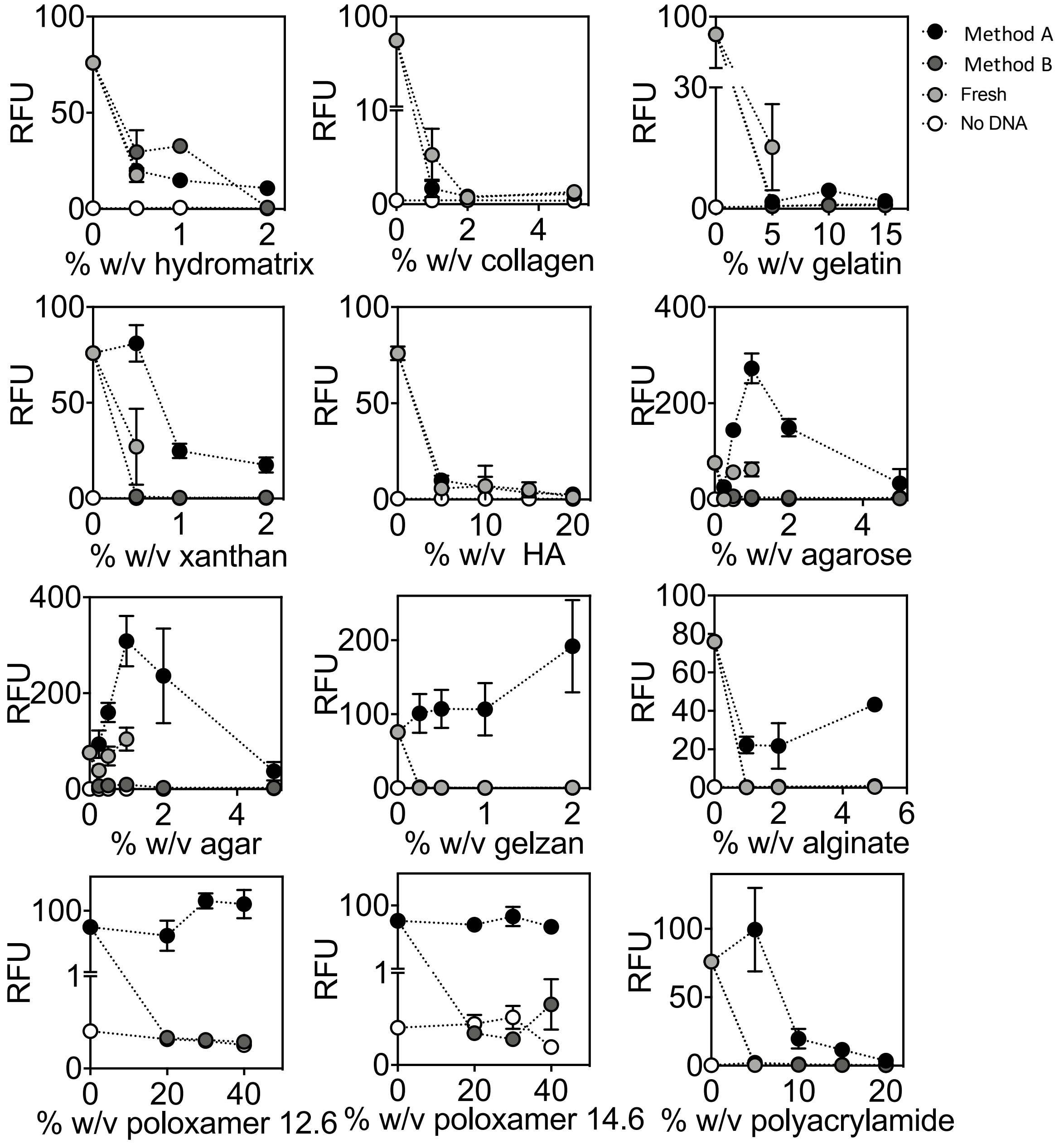


**Fig. S2** Cell-free protein synthesis (CFPS) of mCherry in hydrogels prepared either fresh, from freeze-dried hydrogels (Method A) or with freeze-dried cell-free reagents (Method B) ($n$ = 3, error bars are standard error (SE) mean). RFU = relative fluorescence units. The maximum average RFU over a 16 h incubation is plotted.

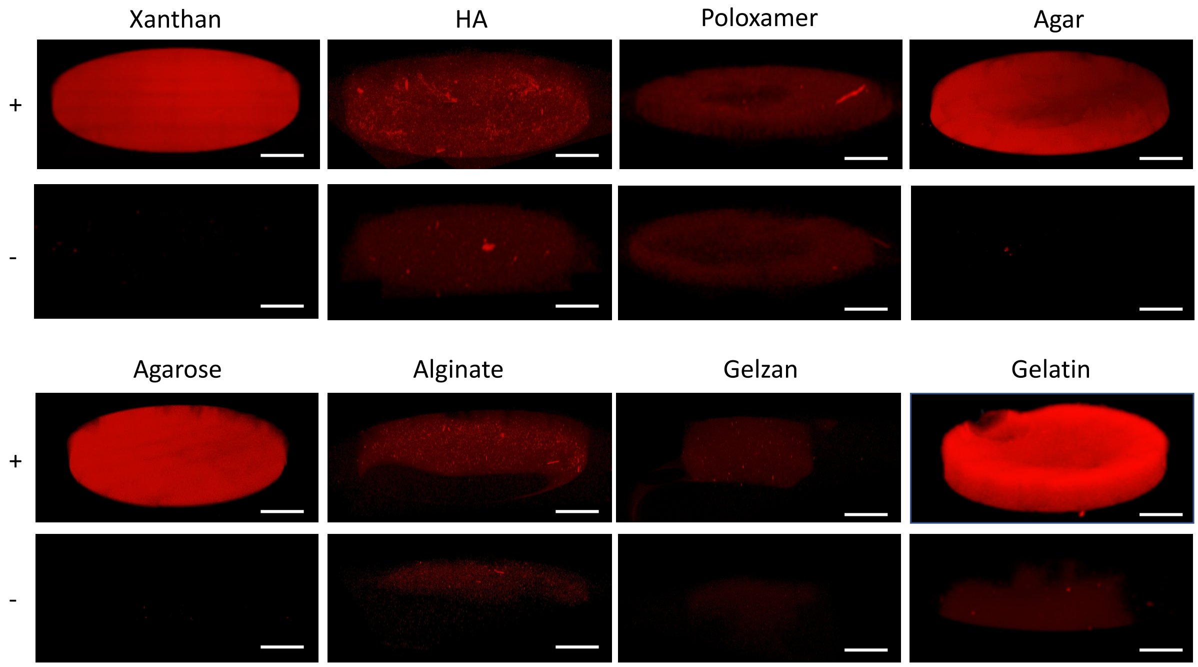


**Fig. S3** Confocal microscopy of hydrogels prepared with cell-free reagents in the presence of (top row) and absence of (bottom row) the mCherry template after 4 h of incubation at 37 °C.

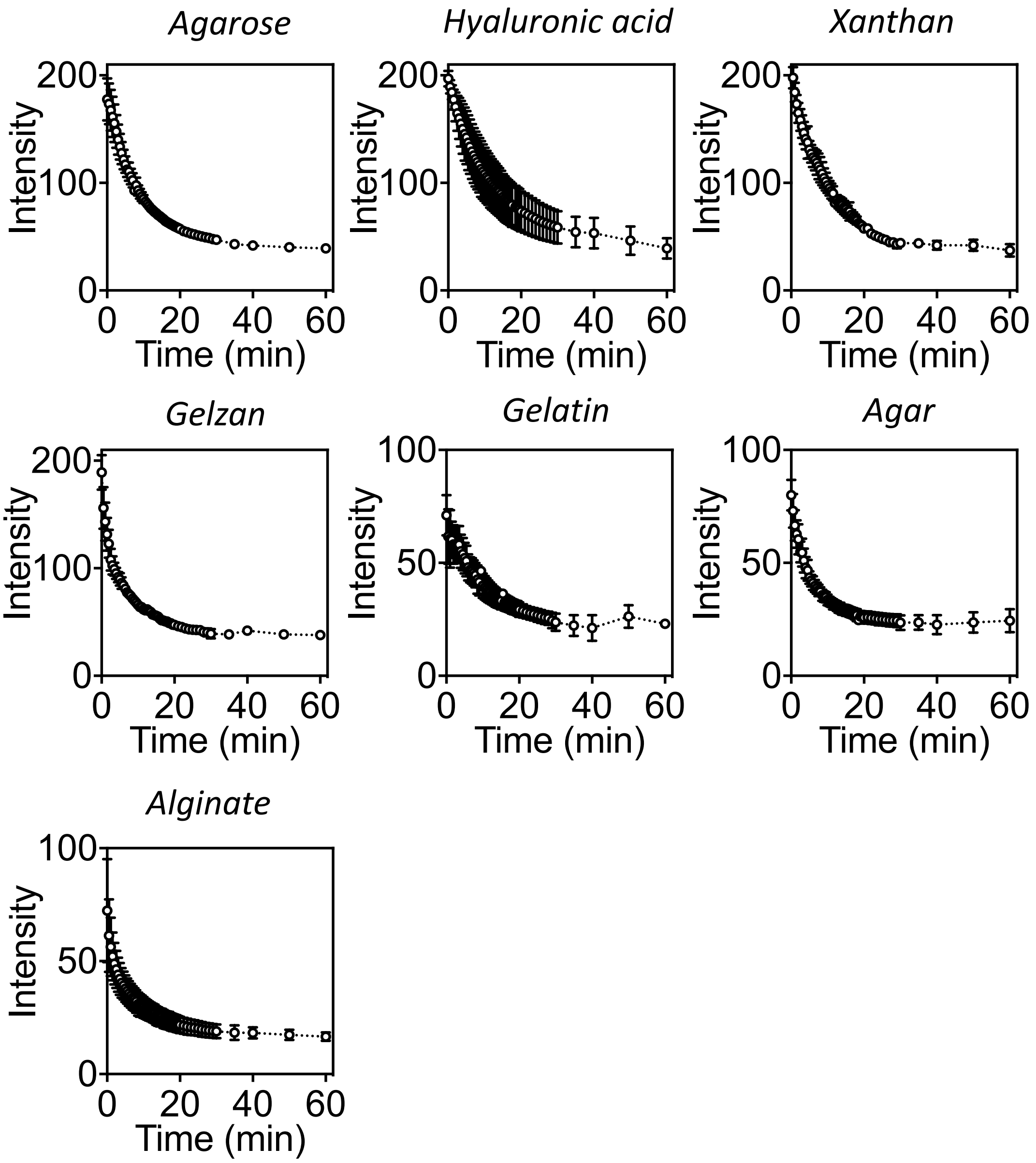


**Fig. S4** The intensity of fluorescein in hydrogels to observe diffusion from a single point over time ($n$ = 3, error bars are SE mean).

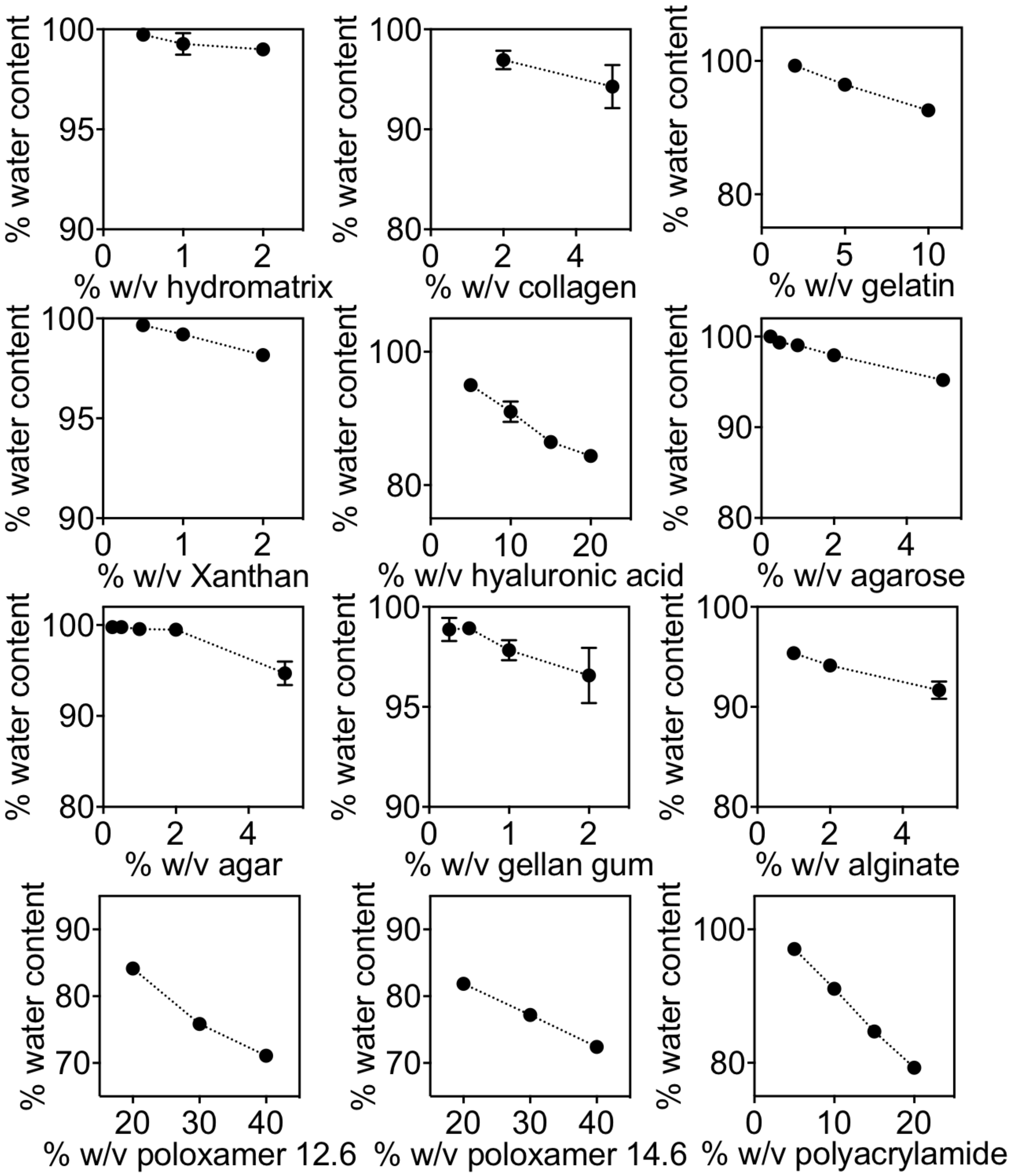


**Fig. S5** Percentage water content of each hydrogel at varying % $^{w}/_{v}$ ratios ($n$ = 3, error bars are SE mean).

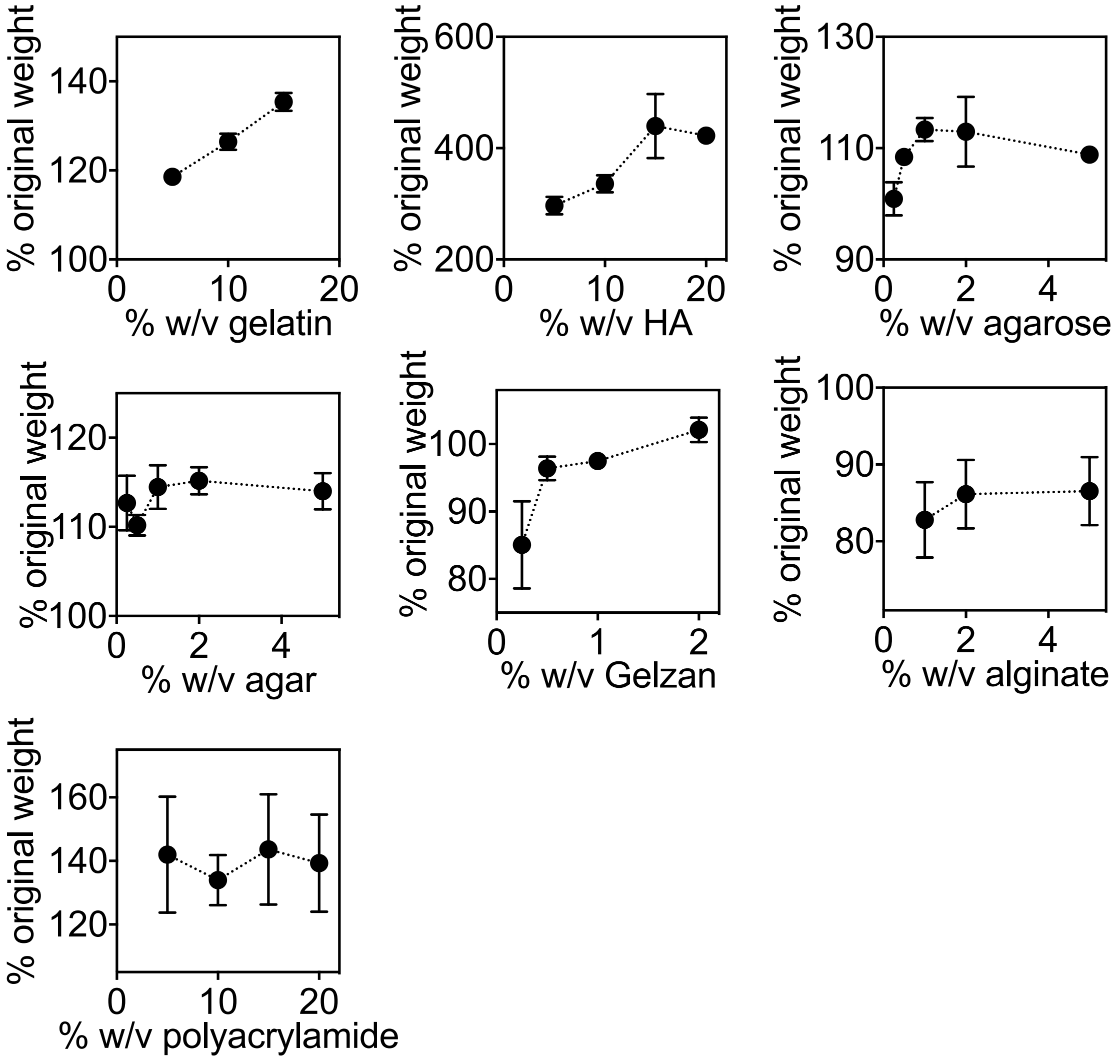


**Fig. S6** Swelling ability of each hydrogel at varying % $^{w}/_{v}$ ratios. Data is shown as a % of the original weight (*n* = 3, error bars are SE mean).

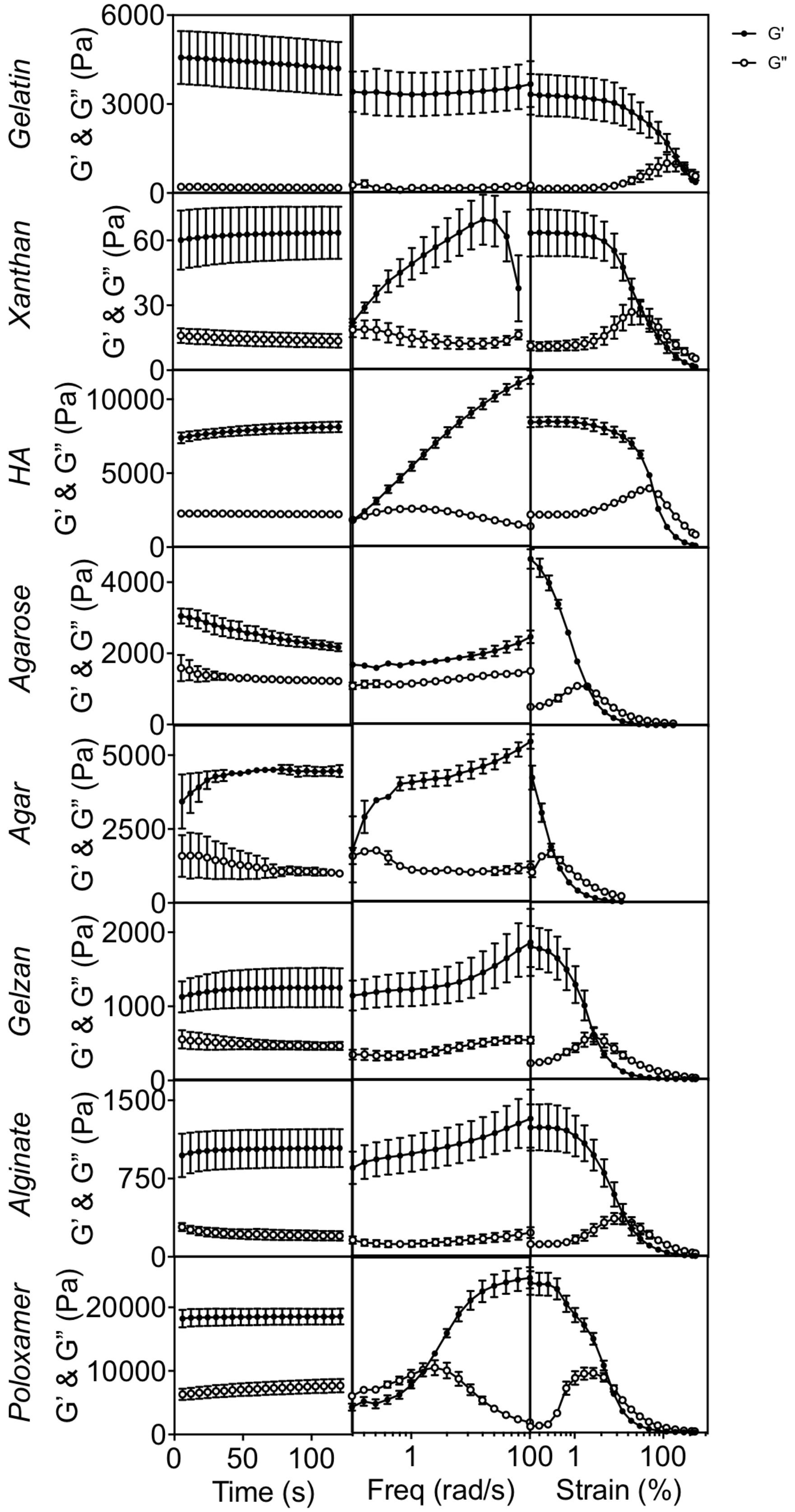


**Fig. S7** Rheological analysis of each hydrogel monitoring the storage modulus (G') and the loss modulus (G") over time (left), over a frequency sweep (central) and a strain sweep (right) ($n$ = 3, error bars are SE mean).

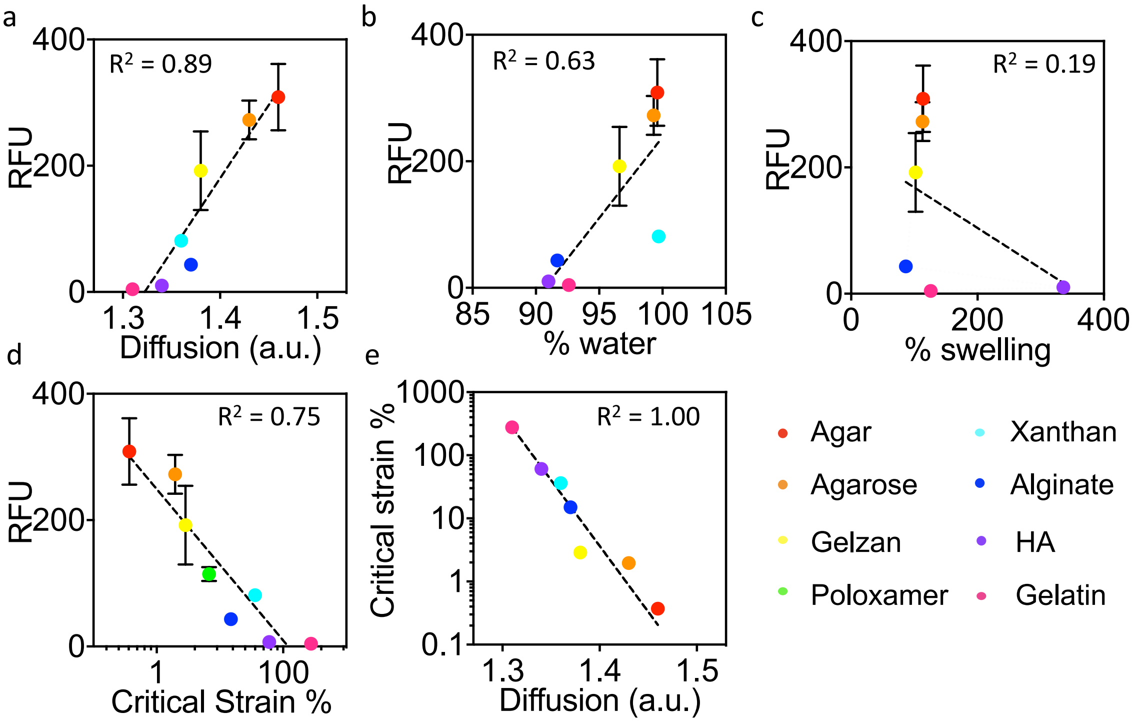


**Fig. S8** The correlation between material properties and the max RFU of mCherry in several hydrogels, (**a**) diffusion (diffusion is represented as a change in fluorescein intensity over time), (**b**) water content, (**c**) swelling, (**d**) critical strain and (**e**) the comparison between the critical strain and diffusion of each material (*n* = 3, error bars are SE mean).

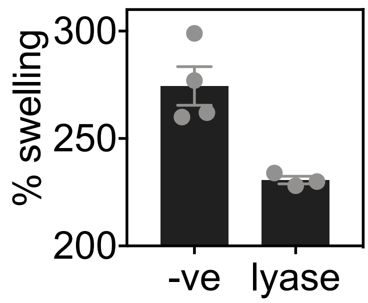


**Fig. S9** The swelling ability of HA hydrogels after 4 h of incubation without and in the presence of the lyase coding plasmid. % swelling is reported as a comparison to the original gel weight (*n* = 6, error bars are SE mean).

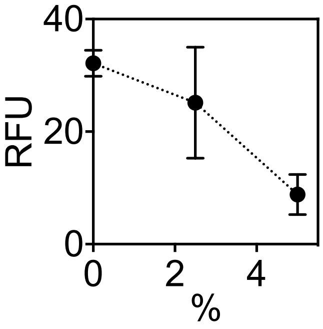


**Fig. S10** CFPS of mCherry in HA:BDDE hydrogels (*n* = 3, error bars are SE mean). RFU is reported as the maximum over a 16 h incubation and % is the % $^{w}/_{v}$ of HA:BDDE.

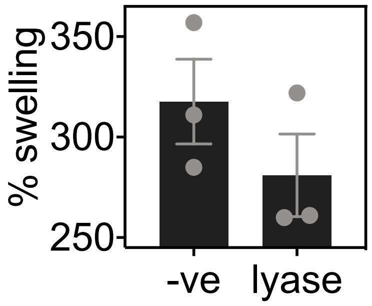


**Fig. S11** The swelling ability of HA:BDDE hydrogels after 4 hours of incubation without and in the presence of the lyase coding plasmid. % swelling is reported as a comparison to the original gel weight (*n* = 5, error bars are SE mean).

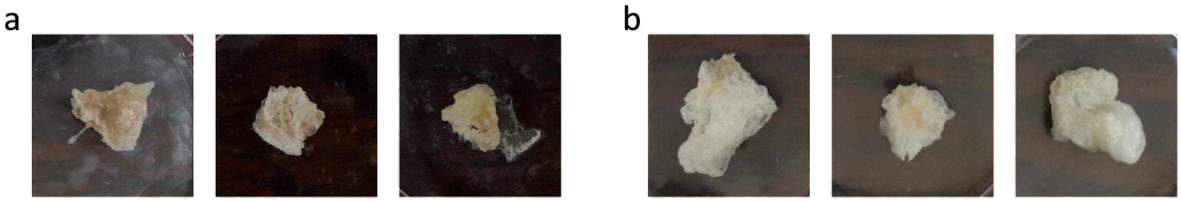

**Fig. S12** Images of HA:BDDE hydrogels prepared with CF components with (**a**) and without (**b**) the lyase coding plasmid after incubation at 37 °C.

### Supplementary Tables

**Table S1** Summary of the hydrogels trialled in this study.

| Material class | Material | CFPS Compatibility | Method A compatible | Method B compatible |
|---|---|---|---|---|
| Peptide | collagen | X | X | |
| | gelatin | X | X | |
| | HydroMatrix™ | X | X | X |
| Polysaccharide | alginate | X | X | |
| | agar | X | X | X |
| | Xanthan gum | X | X | X |
| | Gelzan™ | X | X | |
| | hyaluronic acid | X | X | |
| | agarose | X | X | X |
| Micelle | poloxamer | X | X | |
| Covalent | polyacrylamide | X | X | |

X = CF system performs better in the hydrogel than the aqueous system (hydrogel > aqueous)

X = CF system works but less so than the aqueous system (90 % < aqueous)

**Table S2** Water content, swelling, diffusion and photos of the hydrogels. Data is described as the mean ± SE mean (*n* = 3) to 3 s.f. All values are a percentage.

| Type | Material | % w/v | Water content | Swelling | Reconstitution | Photo |
|---|---|---|---|---|---|---|
| Unstructured | Hyaluronic acid | 5.0 | 95.0 ± 1.00 | 297 ± 9.07 | 100 ± 0.00 | 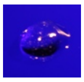 |
| | | 10 | 91.0 ± 1.53 | 336 ± 8.74 | 100 ± 0.00 | 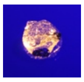 |
| | | 15 | 86.5 ± 0.296 | 440 ± 33.2 | 100 ± 0.00 | 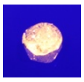 |
| | | 20 | 84.0 ± 0.371 | 423 ± 7.22 | 100 ± 0.00 | 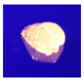 |
| | Collagen | 1.0 | - | - | | |
| | | 2.0 | 96.9 ± 0.917 | - | 81.8 ± 26.8 | |
| | | 5.0 | 94.3 ± 2.17 | - | 120.9 ± 17.9 | |
| | HydroMatrix ™ | 0.5 | 99.7 ± 0.176 | - | 100 ± 0.00 | |
| | | 1.0 | 99.3 ± 0.546 | - | 100 ± 0.00 | |
| | | 2.0 | 99.0 ± 0.000 | - | 100 ± 0.00 | |
| | Alginate | 1.0 | 95.4 ± 0.694 | -17.2 ± 2.84 | 36.4 ± 2.24 | 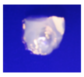 |
| | | 2.0 | 94.1 ± 0.481 | -13.9 ± 2.57 | 28.6 ± 5.02 | 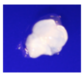 |
| | | 5.0 | 91.7 ± 0.865 | -13.5 ± 2.56 | 28.2 ± 2.32 | 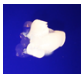 |
| | Agarose | 0.25 | 100 ± 0.00 | 0.900 ± 2.10 | 26.1 ± 0.484 | 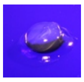 |
| | | 0.50 | 99.3 ± 0.333 | 8.43 ± 0.0670 | 33.7 ± 1.52 | 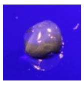 |
| | | 1.0 | 99.0 ± 0.524 | 13.3 ± 1.20 | 40.0 ± 2.96 | 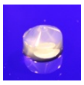 |
| | | 2.0 | 97.9 ± 0.0333 | 13.0 ± 3.63 | 49.3 ± 0.593 | 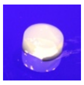 |

| | | | | | | |
|---|---|---|---|---|---|---|
| | | 5.0 | 95.2 ± 0.153 | 8.83 ± 0.524 | 41.8 ± 5.21 | 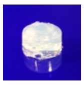 |
| | Agar | 0.25 | 99.8 ± 0.418 | 12.7 ± 3.05 | 24.1 ± 3.18 | 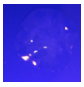 |
| | | 0.50 | 99.8 ± 0.689 | 10.2 ± 1.17 | 36.1 ± 2.63 | 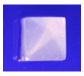 |
| | | 1.0 | 99.6 ± 0.219 | 14.5 ± 2.44 | 40.7 ± 3.07 | 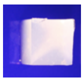 |
| | | 2.0 | 99.5 ± 0.669 | 15.2 ± 1.53 | 50.1 ± 5.09 | 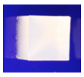 |
| | | 5.0 | 94.7 ± 1.30 | 14.0 ± 2.04 | 61.0 ± 8.68 | 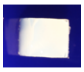 |
| | Xanthan Gum | 0.5 | 99.7 ± 0.133 | - | 100 ± 0.00 | 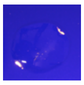 |
| | | 1.0 | 99.2 ± 0.252 | - | 100 ± 0.00 | 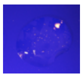 |
| | | 2.0 | 98.2 ± 0.285 | - | 100 ± 0.00 | 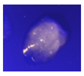 |
| | Gelatin | 2.0 | 99.3 ± 0.200 | 18.6 ± 0.841 | 100 ± 0.00 | 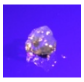 |
| | | 5.0 | 96.4 ± 0.353 | 26.5 ± 1.05 | 39.5 ± 5.83 | 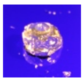 |
| | | 10 | 92.6 ± 0.351 | 35.4 ± 1.16 | 57.8 ± 12.1 | 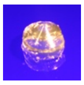 |
| | Gelzan™ | 0.25 | 98.9 ± 0.578 | -14.9 ± 6.46 | 50.4 ± 5.89 | 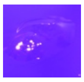 |
| | | 0.50 | 98.9 ± 0.260 | -3.60 ± 1.77 | 47.8 ± 4.20 | 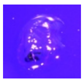 |
| | | 1.0 | 97.8 ± 0.498 | -2.50 ± 0.503 | 65.4 ± 8.48 | 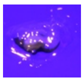 |
| | | 2.0 | 96.6 ± 1.38 | 2.10 ± 1.80 | 100 ± 0.00 | 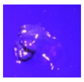 |
| Micelle | Pluronic acid F108 | 20 | 84.2 ± 0.722 | - | 29.2 ± 1.87 | 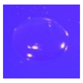 |
| | | 30 | 75.8 ± 0.260 | - | 42.2 ± 4.69 | 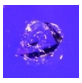 |

| | | | | | | |
|---|---|---|---|---|---|---|
| | | 40 | 71.1 ± 0.252 | - | 45.5 ± 0.00 | 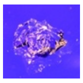 |
| | Pluronic acid F127 | 20 | 81. 9 ± 0.0667 | - | 35.9 ± 2.00 | 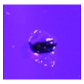 |
| | | 30 | 77.2 ± 0.153 | - | 42.1 ± 0.612 | 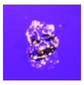 |
| | | 40 | 72.4 ± 0.493 | - | 47.6 ± 2.42 | 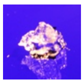 |
| Structured | Polyacrylamide | 5.0 | 97.1 ± 0.133 | 142 ± 10.5 | 30.4 ± 3.76 | 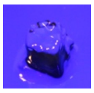 |
| | | 10 | 91.1 ± 0.152 | 134 ± 4.58 | 48.7 ± 5.04 | 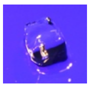 |
| | | 15 | 84.7 ± 0.458 | 144 ± 10.0 | 57.0 ± 2.73 | 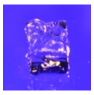 |
| | | 20 | 79.3 ± 0.145 | 139 ± 8.84 | 67.1 ± 0.524 | 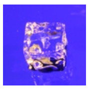 |

**Table S3** Rheological characterisation of hydrogels. Data is described as the mean ± SE mean (*n* = 3) to 3 s.f.

| Hydrogel | G' (Pa) | G" (Pa) | Critical Strain (%) |
|---|---|---|---|
| Agar | 4460 ± 170 | 1040 ± 107 | 0.365 ± 0.0150 |
| Agarose | 2250 ± 175 | 1240 ± 36.4 | 2.00 ± 0.0432 |
| Gelzan™ | 1250 ± 374 | 467 ± 75.2 | 3.01 ± 0.340 |
| Poloxamer | 18500 ± 1780 | 7480 ± 1480 | 6.72 ± 0.0377 |
| Xanthan | 63.4 ± 17.2 | 13.8 ± 4.48 | 39.6 ± 6.43 |
| Alginate | 1040 ± 259 | 203 ± 64.9 | 14.8 ± 2.81 |
| Hyaluronic acid | 8050 ± 484 | 2240 ± 18.0 | 60.5 ± 1.79 |
| Gelatin | 4250 ± 1270 | 179 ± 15.5 | 291 ± 64.5 |

**Table S4** Oligonucleotides used in this study

| Amplification region | Forward primer (5’-3’) | Reverse primer (5’-3’) |
|---|---|---|
| pTU1-A- | TAGAGTCACACTGGCTCACC TTCGG | GCCAGCTGCATTAATGAATC GGCCAA |
| Sequencing GOI | GGCGTATCACGAGGCAGAAT TTCAGATA | TTTGAGTGAGCTGATACCGC TCGC |

**Table S5** Constructs used in this study. For the toehold system, a toehold and trigger (toehold and trigger 1) were selected from Green *et al*. (1) and modified for inclusion as synthetic gene fragments (IDT) in the pTU1_A_ EcoFlex backbone. All other sequences are unmodified from the original citations with the exception that EcoFlex compatible ends were included for cloning.

| | Backbone | Promoter | RBS | Gene | Terminator |
|---|---|---|---|---|---|
| mCherry (2) | pTU1-A_ | J23100 | pET-RBS | *mCherry* | Bba_B0015 |
| eGFP (2) | pTU1-A_ | J23100 | pET-RBS | *eGFP* | Bba_B0015 |
| LacR repressor (3) | pTU1-A_ | J23100 | pET-RBS | *LacR* | Bba_B0015 |
| LacR response (3) | pTU1-A_ | trc | RBS2 | *mCherry* | Bba_B0015 |
| Toehold (1) | pTU1-A_ | Toehold-RBS1 | | *GFP* | Bba_B0015 |
| Trigger (1) | pTU1-A_ | Trigger | | | Bba_B0015 |
| Lyase (4) | pTU1-A_ | J23100 | pET-RBS | *alyA* | Bba_B0015 |

**Table S6** Hydrogel preparation methods

| Hydrogel | Method |
|---|---|
| Hyaluronic acid | The powder was weighed into 1.5 mL micro centrifuge tubes and made up to the required volume with $H_2O$. The gel was allowed to equilibrate at room temperature for 1 h before weighing either 50 mg of the gel directly into the 384-well plate or 500 mg into the Petri dish. |
| Collagen | For CFPS reactions, collagen was measured directly into the 384-well plate. Solution was added directed to the solid and allowed to equilibrate for 30 min at room temperature. For characterisation, the material was weighed directly into a Petri dish and made by the slow addition of solution and mixing using a spatula. |
| Agarose | The powder was weighed into 1.5 mL micro centrifuge tubes and made up to 1 mL with $H_2O$. The tubes were heated to 95 °C for 10 min before cooling to 60 °C to pipette. |
| Agar and Gelatin | In each instance, the powder was weighed directly into a glass conical flask, followed by the addition of water. The mixture was microwaved to dissolve and allowed to cool to 45 °C before pipetting into the 384-well plate or Petri dish. |
| Gelzan | Gelzan was prepared by the addition of the required material powder to a conical flask followed by the addition of $H_2O$ and heating to 80 °C and stirring to dissolve. Once dissolved, the solution was pipetted into either the CF reaction mixture or 6 mM $MgSO_4$ (the concentration of $MgSO_4$ found in the CF reaction). |
| Alginate | Alginate was prepared by weighing the powder into a conical flask and adding $H_2O$. The mixture was stirred until dissolved. 50 µL for CF reactions or 200 µL for material analysis of CF components |

| | |
|---|---|
| | containing 50 mM $CaCl_2$ or purely 50 mM $CaCl_2$ was added to a 1.5 mL microcentrifuge tube. Sodium alginate solution was added in a 1:1 $^{v}/_{v}$ ratio to the tube by pipetting directly into the $CaCl_2$. After 30 min, the gel was removed using a spatula are transferred to either a 384-well plate or vessel for analysis. |
| F-108 (mw ≈ 14600 g/mol) and F-127 (mw ≈ 12600 g/mol) pluronic acid | In each instance, the polymer powder was weighed into a glass vial and the $H_2O$ was added directly. Vials were cooled to 4 °C and stirred to aid dissolving. Solution was then pipetted to the appropriate vessel for analysis and allowed to heat to room temperature to form a gel. |
| Polyacrylamide | Polyacrylamide gels were prepared following the ratios on Supplementary Table 7. Gels were prepared in 2.0 mL microcentrifuge tubes and pipetted to the appropriate vessel immediately are the reagents are added and then allowed to polymerise for at least 60 min. Gels were then transferred to dialysis tubing and dialysed for a minimum of 2 h in $H_2O$. The hydrogel was freeze-dried for future use. |
| HA:BDDE | Hyaluronic acid (HA): 1,4-Butanediol diglycidyl ether (BDDE) hydrogels were prepared based on the synthesis described by Kenne *et al* (5). Briefly, 10 % HA hydrogels were prepared by weighing 100 mg HA into a glass vial, followed by the addition of 1 mL 0.25 M NaOH containing 40 µL (0.216 mmol) BDDE. The mixture was stirred at 35 °C for two h. The gel was then transferred to dialysis tubing and dialysed in $H_2O$ for 48 h. The hydrogel was freeze-dried for future use. |

**Table S7** Polyacrylamide gel preparation

| | Initial concentration | Vol (μL) | | | | Final concentration |
|---|---|---|---|---|---|---|
| | | 5 % | 10 % | 15 % | 20 % | |
| 30 % acrylamide/ bis acrylamide | 30 | 266.7 | 533.3 | 800.0 | 1065.6 | 5, 10, 15, 20 |
| Tris, pH 7.5 | 10 mM | 480 | | | | 30 μM |
| TEMED | 66.9 mM | 32 | | | | 1.34 mM |
| APS | 10 % | 8 | | | | 0.05 % |
| $H_2O$ | - | 813.4 | 545.6 | 280.0 | 14.4 | - |
| Total | | 1600 | | | | |
